# A novel green approach for treatment of immature Schistosomiasis Mansoni infection in mice; Arabic gum (Acacia Senegal) antischistosomal properties

**DOI:** 10.1101/347278

**Authors:** Rabab Selem, Samia Rashed, Mohammad Younis, Boshra Hussien, Fatma Mohamed, Awatif Edrees, Asmaa EL-kholy, Gehan Rashed, Shereen Kishik, Ahlam Moharm, Marwa Nageeb, Manal Kardoush

## Abstract

Schistosomiasis is one of the most socioeconomically exhausting parasitic infection in tropical and subtropical areas. Praziquantel (PZQ), the only common schistosocidal drug in use, is not efficient enough for treatment of immature infection. Arabic gum (AG) is a complex polysaccharide acts as anti-oxidant which modulates the inflammatory and/or immunological processes. This study explores for the first time, the antischistosomal properties of AG in mice infected with the immature stage of *Schistosoma mansoni*. Mice were divided into four groups: control group (infected non-treated), AG treated group, PZQ treated group, and AG+PZQ treated group. Oral administration of AG in a dose of 1gm/kg body weight, daily for 3 consecutive weeks post-infection (p.i.) resulted in a statistically significant lower worm burden in both AG group and AG+PZQ group compared to PZQ and control groups. AG+PZQ group always showed the best performance when compared with other groups regarding tissue egg load and oogram pattern. AG, both alone and in combination with PZQ, decreased the number, diameter; increased the cellularity and the number of degenerated Schistosoma eggs inside granulomas. Results obtained by this work elucidated a promising AG bioactivity against *S. mansoni* immature stages and provided a platform for subsequent experimental studies to illuminate the academia more about this novel and “green” antischistosomal agent.

**Author summary:** Schistosomiasis is a major public health threat in many parts of the world, it affects more than 240 million people in more than 70 countries and almost 800 million people are at risk of acquiring this disease. Serious consequences and disabilities might result from untreated schistosomiasis such as hepatosplenic fibrosis with portal hypertension, gastrointestinal hemorrhage and death.

Schistosomiasis control is focused on periodic treatment with praziquantel (PZQ). However, PZQ has only moderate action against young developing stages of schistosomula. Recently, resistance has emerged to PZQ. Therefore, chemotherapy alone is unlikely to reduce infection levels of schistosomiasis. Several practical approaches have been suggested to augment treatment programs. Of course, the development of a complementary treatment would contribute enormously to the reduction of schistosomiasis. Recently, natural products have been popular and attracted most of the attention as it could offer new effective therapy against schistosomiasis. Arabic gum (AG) is an edible, dried sticky exudate from *Acacia Senegal*, which is used in this study to assess the AG antischistosomal properties. Our study revealed that AG has an excellent statistically significant effect against immature murine schistosomiasis, both alone and in combination with PZQ. This approach may point to novel targets for treatment of schistosomiasis.

## Introduction

Schistosomiasis is the most common disease caused by parasitic worms, known as blood flukes, it affects over 240 million people around the world with almost 800 million are at risk of infection [1]. Serious consequences and disabilities might result from untreated schistosomiasis such as chronic malnutrition, anemia, organ scarring and fibrosis [2].

Control of such long-term morbidity is a priority of the World Health Organization (WHO), it adapts a preventive strategy via mass drug administration campaigns [3]. Praziquantel is the drug of choice for treating all species of Schistosoma. Unfortunately, some strains have developed a resistance against it making their treatment a challenge [4-7]. Although praziquantel is highly effective against adult schistosomes and very early stage of schistosomula just few hours after penetration into the host’s skin, it is much less effective against young developing stages of schistosomula [8], Thus, it is essential to develop a new irresistible alternative lacking the aforementioned drawbacks [9].

Arabic gum (AG) is a dried exudate obtained from stems and branches of *Acacia senegal* (Leguminosae), consisting of calcium, magnesium, and potassium salts of the polysaccharide Arabic gum acid [10]. It has been used in Arabic folk medicine to treat patients suffering from chronic renal failure as it decreases the requirements as well as the frequency of hemodialysis [11]. US Food and Drug Administration have listed AG as one of the safest dietary fibers [12].

Different studies showed that AG can modulate TGF-b1 generation and function [13], stimulated mouse dendritic cells [14], control chemical plaque in subjects with gingivitis [15], and exert a cytoprotection against Hg-induced nephrotoxicity [16].

Other studies reported several favorable renal effects including reduced plasma phosphate concentration, blood pressure, proteinuria, as well as extra renal effects such as slowing of intestinal glucose transport, which might be of value in the prevention and treatment of obesity and diabetes [17, 18]. It has been also reported to induce fetal hemoglobin in sickle cell anemia [ 19], prevents and enhances healing of gastric ulcers [20], influences the expression of murine ovarian oxidative stress gene [21] and improves semen quality and oxidative stress capacity in alloxan-induced diabetes in rats [22].

As antimicrobial agent, AG was reported to be an efficient antimicrobial agent, of a natural origin, against many buccal microorganisms such as *Prophyromonas gingivalis* and *Prevotella intermedia* [23] fungi as *Candida albicans*, *Aspergillus niger* and *Microsporum canis* [24] bacteria as *Staphylococcus aureus, Staphylococcus epidermidis, Streptococcus pneumoniae, Salmonella typhi, Ps. aeroginosa*, [25]. As far as we know, only one published parasitological study has investigated the antimalarial effect of AG, it stated that AG slightly, yet, significantly decreased the parasitaemia and significantly expanded the life span of the infected mice [26].

The aim of this study was to explore and evaluate the antischistosomal properties of AG in mice infected with *Schistosoma mansoni* at the immature stage.

## Materials and methods

A pilot study was done on about 30 mice 2 months before the main experiment to assess if there is any antischistosomal activity of AG which has been given as 10% in the drinking bottles for 3 weeks starting from the day of infection. The results of AG on the scale of total worm load was remarkable and encouraging to lunch the main experiment.

### Parasites and animals

Fifty laboratory-bred male Swiss albino mice, CD1 bred, were used in this study, as this was the minimum required number to guarantee statistically reliable and reproducible results. Cercariae of *S. mansoni* were obtained from infected *Biomphalaria alexandrina* snails, which were reared and maintained at Schistosome Biological Supply Program (SBSP), Theodor Bilharz Research Institute (TBRI), Giza, Egypt. Each mouse was infected with 80 *S. mansoni* cercariae suspended in 0.2 ml water via subcutaneous injection [27].

### Ethics Statement

This study was conducted in accordance with legal ethical guidelines of the medical ethics committee of the Theodor Bilharz Research institute (TBRI), Giza, Egypt. Approval no. 4013/2016.

### Experimental design

Mice were divided into 4 groups, 10-13 mice each, representing: AG treated group, PZQ treated group, AG+PZQ treated group and untreated infected control group. AG group mice were treated daily starting from the 1^st^day post-infection (p.i) till the 21^st^ day using a dose of 1 gm/kg body weight (AG is a powdered material obtained from local conventional herbal medicine market, suspended in water as a solvent reagent at a concentration of 100mg/ml). This dose was similar to that of Nasir *et al.* 2012 [ 17], but given individually to each mouse orally using a syringe with a curved end. On the 21^st^& 22^nd^ day PZQ (Alexandria Company for Pharmaceuticals and Chemical Industries, Alex., Egypt) was freshly suspended in 13 ml of 2% cremophore-EL (Sigma Chemical Co., USA) and orally administered to mice at a dose of 500 mg/kg body weight for two consecutive days [28]. Three weeks later (6 weeks p.i) all animals were scarified to assess AG antischistosomal efficacy.

### Evaluation of AG antischistosomal effect

#### 1. Worm Burden

Schistosomes recovery was done by porto-mesenteric perfusion technique, 3 weeks post-treatment, according to the method of Duvall and DeWitt, [29]. Drug efficacy was measured by percent reduction of worms according to the formula of Abdel Salam et al., [30]: *R*% (percent reduction) = *C*–*T/C* ×100, where *C* is the mean worm burdens in control infected animals and *T*, mean number of worms in infected treated animals.

#### 2. Tissue egg load (hepatic and intestinal)

Segments of liver and intestine were blotted between two filter papers, weighed, transferred each to a test tube containing 5 ml 5% potassium hydroxide solution [31], and left overnight at room temperature to facilitate tissue digestion without egg destruction. Next morning, tubes were incubated at 37°C for 1 h to finish the tissue clearance [32]. Ova in homogenous emulsions were counted after being spread on slides, and the number of ova/mg tissues was calculated. To detect the egg load in the hepatic and intestinal tissue, the average number of eggs in 1 ml sample was multiplied by the total volume of potassium hydroxide, then divided by the weight of tissue to yield the number of eggs/gram tissue [33]. Percentage reduction was accordingly calculated. *R*% (percent reduction) = *C*–*T/C* × 100, percentage reduction was calculated using the aforementioned equation [30].

#### 3. Oogram pattern

After mice perfusion, three segments, one cm in length of the small intestine were cut longitudinally, rinsed in saline, slightly dried on filter paper, compressed between two glass slides and examined under microscope for oogram pattern that may reflect the direct drug action on ova development [34]. Eggs of *S. mansoni* were classified in the current study in three types immature, mature and dead [35].

#### 4. Histopathological examination

Mice livers were fixed for 48 h in 10% buffered formalin and then embedded in paraffin. Haematoxylin and eosin were used to stain sections [36] for granuloma counting while Masson trichrome stains [37] were used to demonstrate collagen fibers. Lesions containing single ova in their centers were selected for measurement [38]. The granuloma diameter of each case was measured using the ocular micrometer [39]. For each section, granulomas were counted in five successive fields (10×10).

### Statistical analysis

Gathered data were tabulated and analyzed using SPSS statistical software (IBM Corp., Armonk, NY, USA). Data were expressed as mean ±SD or SE. Analysis of variance between groups was done using ANOVA test and when significant, post hoc Bonferroni test was applied for pairwise comparison between groups. P value <0.05 was considered statistically significant. All statistical tests were two-sided. Chi square test was used to assess if there was a significant difference between granuloma types in various study groups.

## Results

Regarding the worm load, the AG group demonstrated the best reduction rate (75.6%) followed by the AG+PZQ group (72.5%) while the PZQ group had the lowest rate (28.7%). The difference between all groups was statistically significant (P-value <0.001). Comparing each group with the control group, the difference was significant except for the PZQ group (P-value 0.151). While comparing the PZQ group with the AG group and with the AG+PZQ group the difference is statistically significant (P-value 0.002 and 0.008 respectively), Table 1.

**Table 1.**
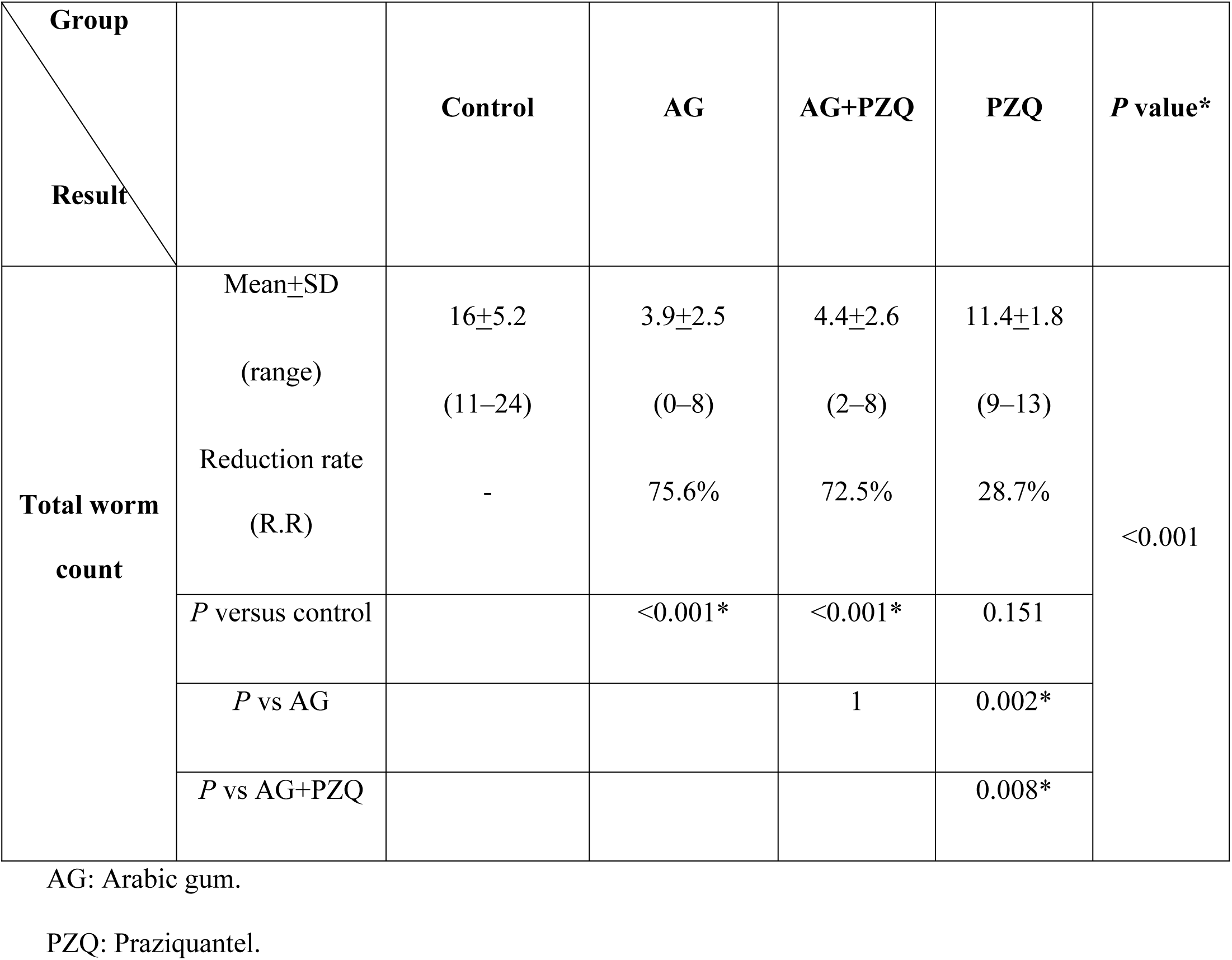
Performance of AG, PZQ and combined AG+PZQ therapeutic regimens on *Schistosoma mansoni* total worm burden after treatment of infected mice during the immature infection stage.

| Group<br>Result |  | Control | AG | AG+PZQ | PZQ | P value* |
| --- | --- | --- | --- | --- | --- | --- |
| Total worm count | Mean±SD | 16±5.2 | 3.9±2.5 | 4.4±2.6 | 11.4±1.8 | <0.001 |
|  | (range) | (11–24) | (0–8) | (2–8) | (9–13) |  |
|  | Reduction rate (R.R) | - | 75.6% | 72.5% | 28.7% |  |
|  | P versus control |  | <0.001* | <0.001* | 0.151 |  |
|  | P vs AG |  |  | 1 | 0.002* |  |
|  | P vs AG+PZQ |  |  |  | 0.008* |  |

AG: Arabic gum.
PZQ: Praziquantel.

About egg count in the liver, the AG+PZQ had the lowest number (950±498.8), followed by the PZQ group (1964.8±909), then the AG group (2315.8±252.7) and the highest number belonged to the control group (8507.4±915.2), with statistically significant difference (P-value <0.001). Comparing each group with the control group, the difference was significant (P-value <0.001). The AG+PZQ group demonstrated the best hepatic egg load reduction rate (88.8 %) followed by the PZQ group (76.9%) while the AG group had the lowest rate (72.7%). Comparing the result of the AG group with that of the AG+PZQ group revealed a statistically significant difference (P-value 0.010) while comparing it to that of the PZQ group revealed a non-significant difference (P-value1). Similarly, comparing the result of the AG+PZQ group to that of the PZQ group revealed a non-significant difference (P-value 0.226).

A similar pattern was noted for the intestinal egg load as the AG+PZQ had the lowest number (961.1±387.2), followed by the PZQ group (1121.8±629), then the AG group (3168.8±1016.7) and the highest number belonged to the control group (7205.1±1049.6), with statistically significant difference (P-value <0.001). Comparing each group with the control group, the difference was significant (P-value <0.001). The AG+PZQ group demonstrated the best intestinal egg load reduction rate (86.6 %) followed by the PZQ group (84.4 %) while the AG group had the lowest rate (56%). Comparing the AG group to either AG+PZQ or PZQ group yielded a statistically significant difference (P-value <0.001 and 0.001 respectively), while comparing the PZQ and AG+PZQ groups yielded a non-significant difference (P-value 1). Table 2.

**Table 2.**
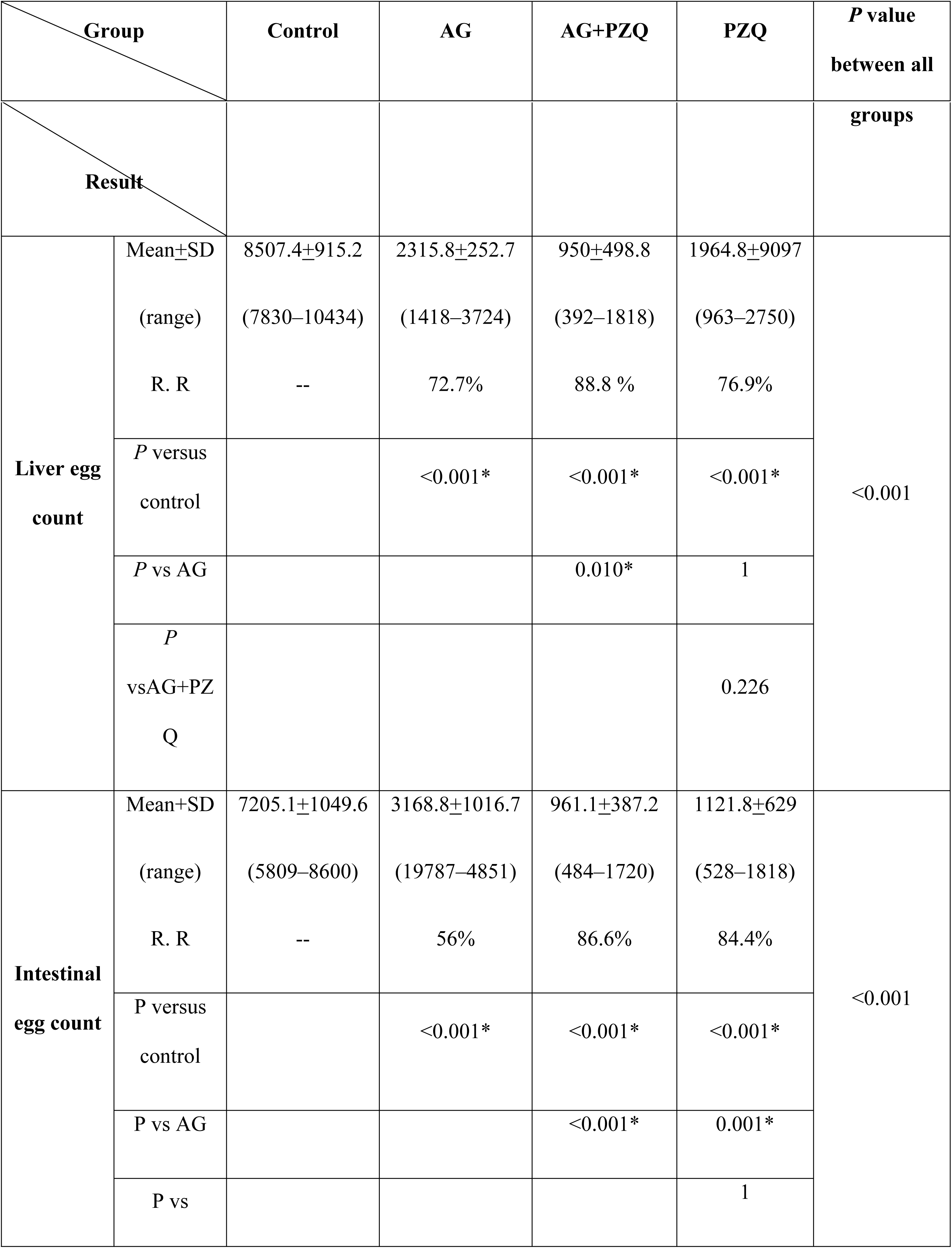
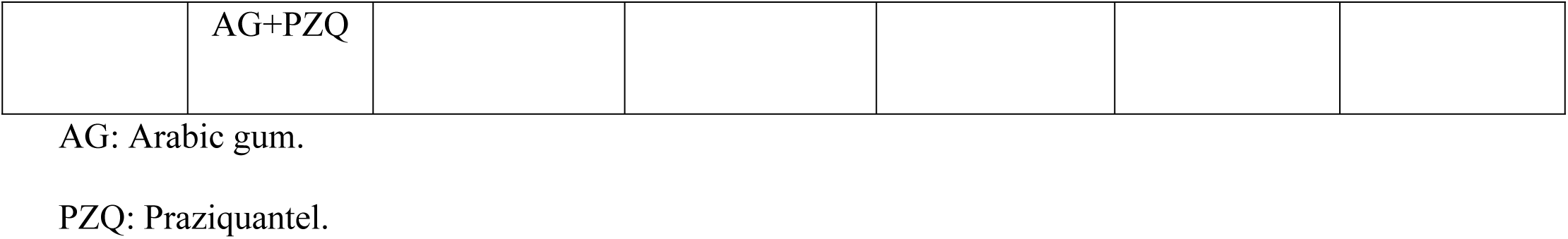
Comparison between the reductive effect of AG, PZQ and combined AG+PZQ therapeutic regimens reductive effect on *Schistosoma mansoni* liver and intestinal egg count after treatment of infected mice during the immature infection stage.

Concerning the oogram pattern, the AG+PZQ presented the best results demonstrating the lowest immature egg count (45±1.7), followed by the PZQ group (51.8±1.8), then the AG group (54.9±6.4) and the highest number belonged to the control group (51.1±4.6), yet the difference was insignificant (P-value 0.015). Comparing each group with the control group, the difference was also insignificant. Comparing the result of the AG group with that of the AG+PZQ group revealed a statistically significant difference (P-value 0.009) while comparing it to that of the PZQ group revealed a non-significant difference (P-value 1). Similarly, comparing the result of the AG+PZQ group to that of the PZQ group revealed a non-significant difference (P-value 0.306).

The mature egg count was (40.4±4.9) in the AG+PZQ group, (42±2.1) in the PZQ group, and (40±6.7) in the AG group. The difference between each group and the control group was statistically insignificant. Comparing the AG group to either AG+PZQ or PZQ group, as well as comparing the AG+PZQ groups yielded non-significant differences (P-value 1, 1 and 1 respectively).

While regarding the dead egg count, the highest number was detected in the AG+PZQ (5–25, Mean +SD14.6±6.8), followed by the control group (4-10, Mean +SD6.3±2.1), then, the PZQ group (5–8, Mean ±SD6.2±1.3). When each group was compared to the control group the difference was statistically insignificant (P-value 1 and 1) except for the AG+PZQ group (P-value 0.001). While comparing the AG group with the AG+PZQ group the difference is statistically significant (P-value <0.001) and statistically insignificant when compared with the PZQ group (P-value 1). On the other side, the difference between the PZQ and the AG+PZQ groups was statistically significant (P-value 0.003), Table 3.

**Table 3.**
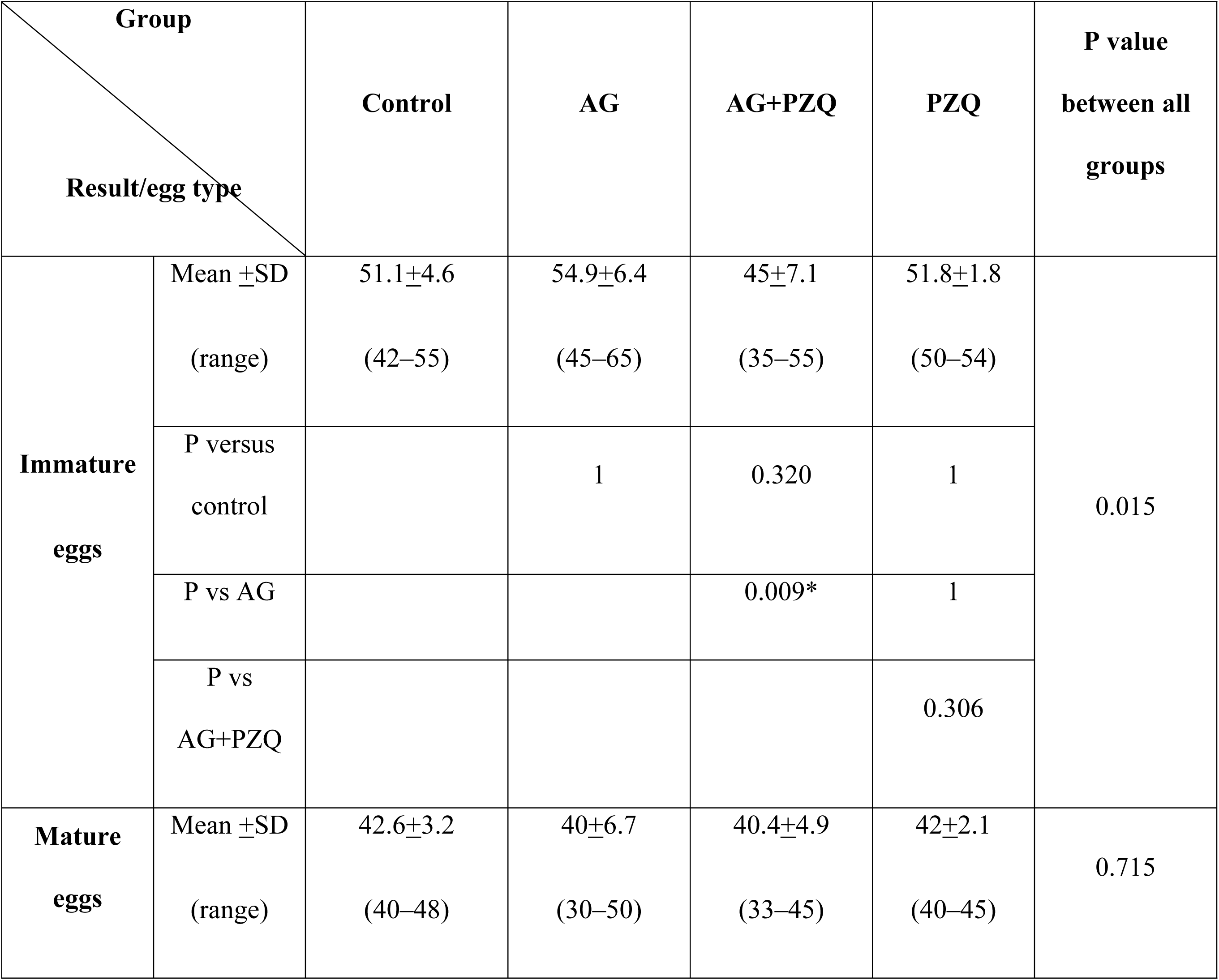
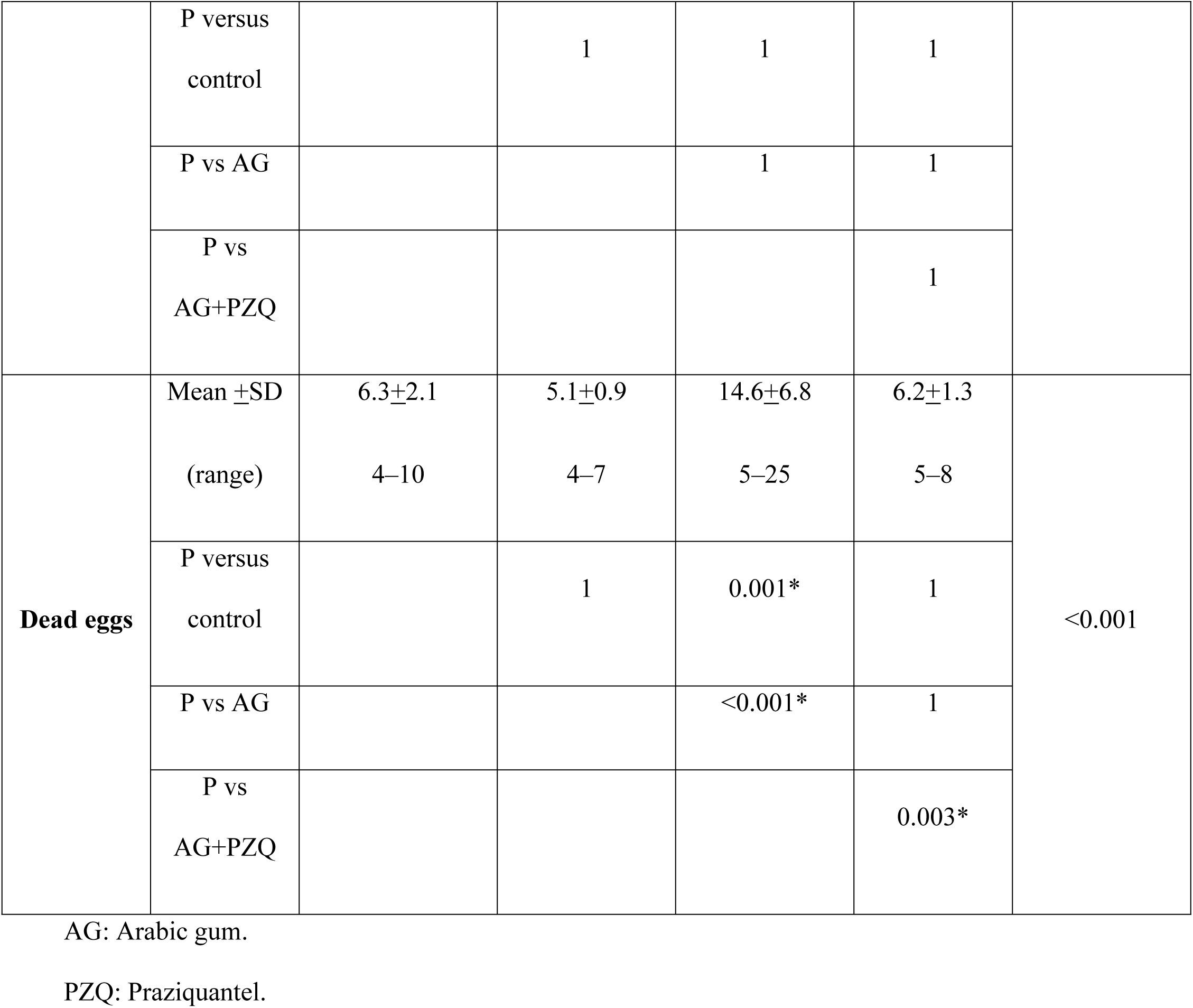
Oogram pattern of AG, PZQ and combined AG+PZQ therapeutic regimens after treatment of infected mice during the immature infection stage.

The histopathological assessment of the granuloma diameter revealed that the smallest diameter belonged to the AG+PZQ (214.23±12.18), followed by the PZQ group (272.22±11.2), then the AG group (297.28±7.5) and the largest diameter belonged to the control group (353.15±12.4). Comparing each group with the control group, the difference was significant (P-value0.0010,0.0010 &0.00010for the AG, PZQ and AG+PZQ groups respectively).

On the other side, the AG+PZQ group demonstrated the lowest granuloma number (3.32±1.21), followed by the AG group (3.9±1.13), then the PZQ group (5.4±1.82), and the control group presented the highest granuloma number (10.62±1.97). Comparing each group with the control group, both the AG and AG+PZQ showed a significant difference (P-value 0.0064&0.0064 respectively) while the difference between the PZQ group and the control group was insignificant (P-value 0.09).

While results of the granuloma type revealed that the AG group had the highest cellular and the least fibro-cellular and fibrous types among all groups (80%,20% &0%), followed by AG+PZQ group (65%, 30%&5%) and the last in order was the PZQ group (55%,43%&2%). Only AG and AG+PZQ had significantly different granuloma types as compared to the control group (P-value <0.001&0.022 respectively), while the types distribution in the PZQ group was insignificantly different than that of the control group (P-value 0.247).

The State of *S. mansoni* eggs demonstrated a different pattern as the lowest number of intact eggs and the highest number of degenerated eggs was detected in the AG group (17 &83 respectively), while the AG+PZQ group had (23) intact eggs and (77) degenerated eggs, and the PZQ group had (45) intact eggs and (55) degenerated eggs. Comparing each group with the control group, the difference was significant (P-value <0.001). Table 4 and Fig 1.

**Fig 1.:**
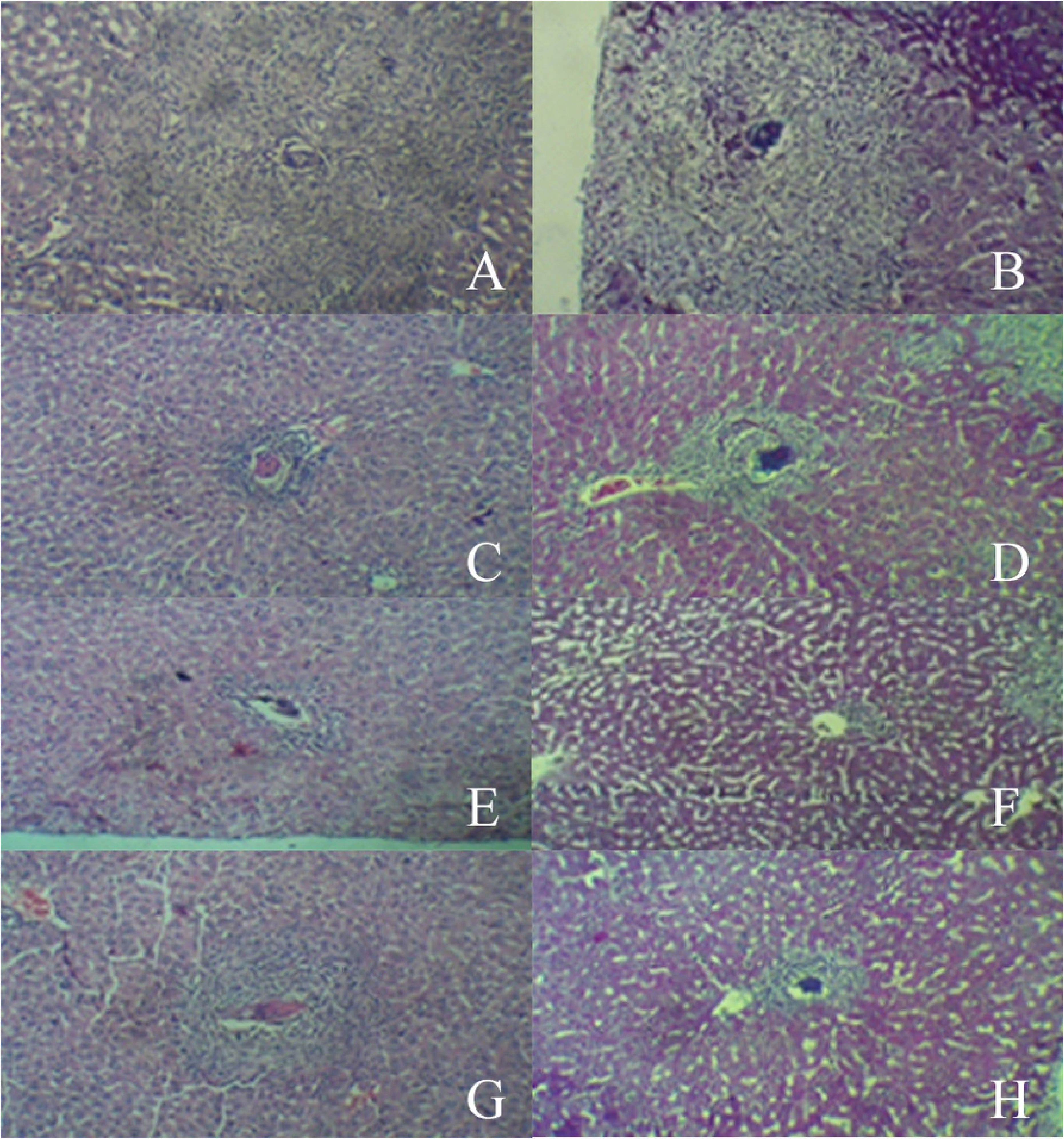
Liver histology at six weeks after *S. mansoni* infection of CD1 bred mice with 80 cercariae by subcutaneous injection (hematoxylin & eosin stain: 100x magnification) 1-A:liver infected sections non treated control mice groups 6 weeks p.i. showing large number of fibrocellular granulomas stained with H&E (x100) 1-B:liver infected sections non treated control mice groups 6 weeks p.i. showing large number of fibrocellular granulomas stained with masson trichrome stain (x100) 1-C: Liver of infected mice group treated PZQ showing less number of fibrocellular granuloma 1-D: Liver of infected mice group treated PZQ showing decrease in granuloma size showing small fibrous granuloma 1-E: Liver of infected mice group treated with AG and PZQ showing less number of cellular granuloma 1-F: Liver of infected mice group treated with AG and PZQ showing decrease in the granuloma size showing small granuloma with degenerated eggs 1-G: Liver of infected mice group treated with AG showing decrease in size of granulomas 1-H: Liver of infected mice group treated with AG showing decrease in size of granulomas degenerated eggs

**Table 4.**
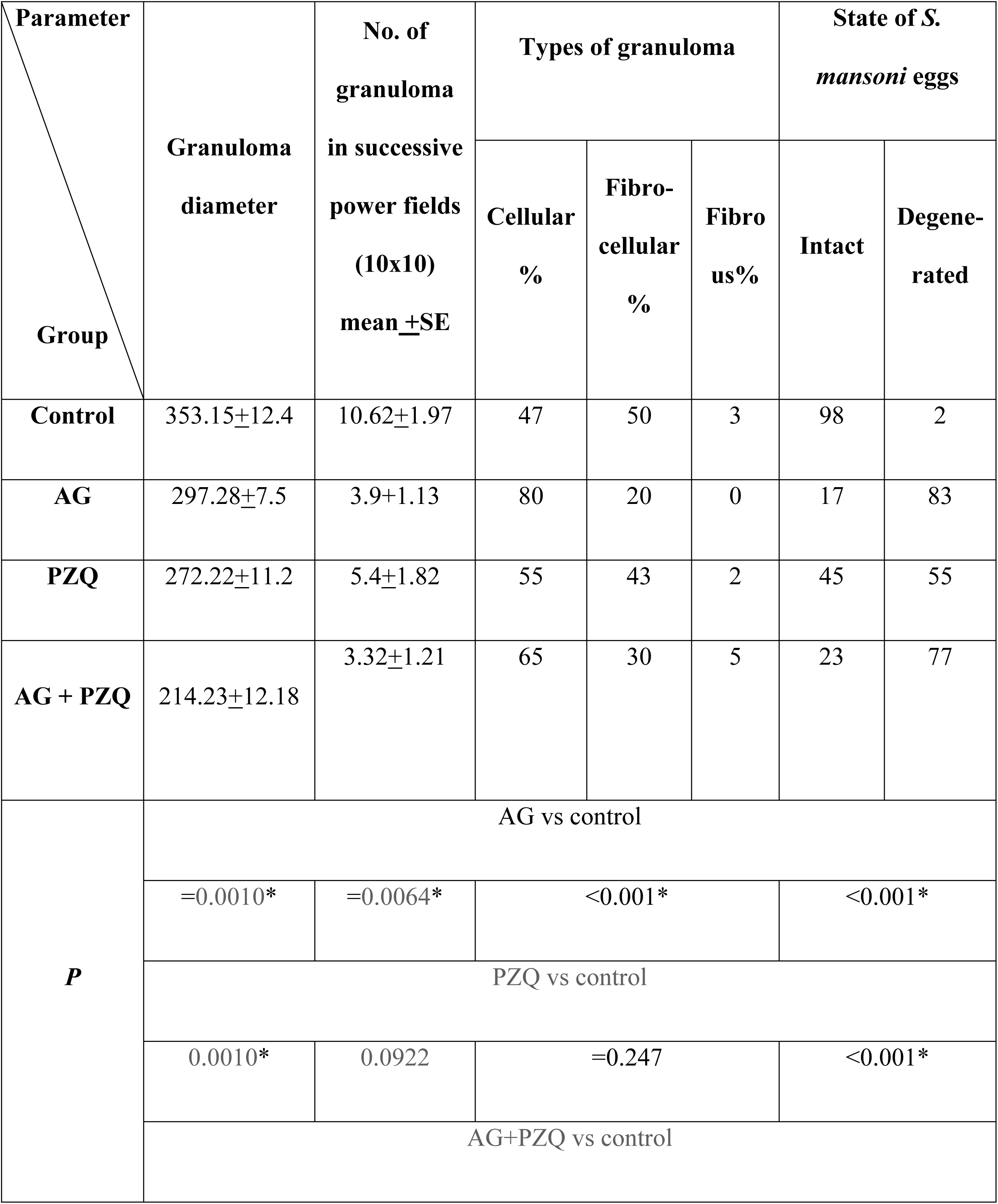
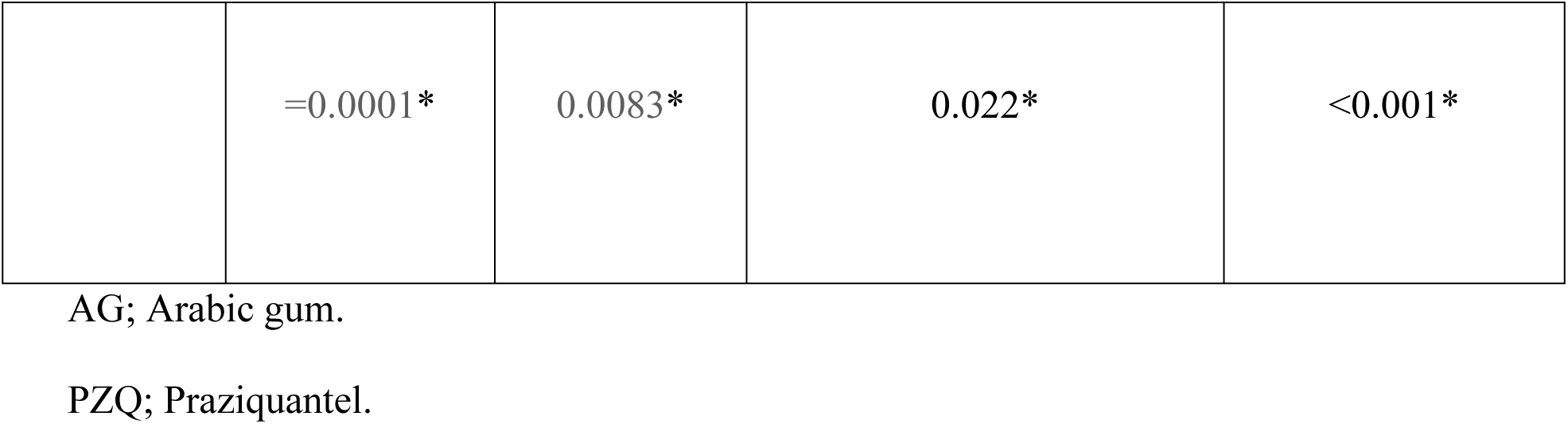
Effect of AG, PZQ and combined AG+PZQ treatment regimens on Schistosoma *mansoni* induced hepatic granulomas parameters as compared with the control group

## Discussion

Schistosomiasis control programs are based mainly on a single drug which is praziquantel tablet [40]. Despite the fact that patients could tolerate PZQ well, it has some drawbacks including the emergence of drug resistance [4, 5], the poor efficacy on the immature stages [41], the large, bitter tablets, and the unavailability of a pediatric formula [42]. Recently, natural products and natural product-derived compounds have been popular and attracted most of the attention as it could offer new effective therapy against schistosomiasis. Arabic gum (AG) is an edible, dried sticky exudate from *Acacia Senegal*, which is rich in soluble dietary fiber [43].

In this study assessment of AG antischistosomal properties revealed an excellent statistically significant effect against immature murine schistosomiasis, both alone and in combination with PZQ demonstrated in parasitological parameters; worm load, egg count, oogram pattern and histopathological results; granuloma metrics (diameter, number. and state of *Schistosoma* eggs within them).

In all parasitological parameters, apart from the worm load, AG+PZQ treated animals showed the best results as compared to monotherapy groups, denoting a considerable synergistic effect of AG+PZQ on both female fecundity, egg maturation and ability to elicit its immunopathological effect. The best reduction rate of *Schistosoma* worms was demonstrated in the AG monotherapy group, nevertheless, the difference between AG and AG+PZQ treated mice worm load was negligible. On the contrary, the PZQ treated mice demonstrated the worst results among all studied groups regarding total worm load, however, such results were expected as PZQ is less effective against immature schistosomiasis, the stage targeted in this experiment.

Regarding the histopathological parameters, the AG+PZQ group showed the least mean granuloma diameter, while the largest diameter was demonstrated in the AG group. This could be explained by the fact that the granuloma of that group is the highest cellular, the least fibrocellular and fibrous granuloma types, lacking adequate fibers amount diminishes its contraction and permits large sizes. Another explanation is based on the highly significant difference in *S. mansoni* intact-degenerated eggs distribution within the examined granulomas, as the cellularity that dominated granulomas of AG treated animals might eliminate the physical barriers which would be created by fibrous tissue and hampers the action of the host immune system. Concerning the mean granuloma number, AG was significantly effective, both alone and in combination with PZQ, followed by the combination of AG+PZQ and the least effect belonged to the PZQ monotherapy. These results could be attributed to the destructive effect of AG on fecundity which in turn decreases the number of evolving granulomas.

The AG therapeutic effect on immature murine schistosomiasis in this experiment could be attributed to its immunomodulatory effect, as it stimulates the dendritic cells [14] which are antigen-presenting cells responsible for triggering both innate and adaptive immunity [44].

Also, it might be attributed to the antioxidant properties of AG in many tissues like renal tissue [16], RBCs in sickle cell anemia (SCA) disease [19] and hepatic tissue as mentioned by Ahmed *et al.,* [45] who stated that AG significantly decreased the level of hepatic enzymes, lipid peroxidation, antioxidant enzymes as well as the expression of oxidative stress genes. Activities of superoxide dismutase (SOD), catalase (CAT) and glutathione peroxidase (GPx), which may contribute to the alleviation of *Schistosoma mansoni* infection consequences similar to what has been reported in many other antioxidants like gold nanoparticles *Ceratonia siliqua* pod extract [46], limonin [47]. Another theoretically potential mechanism of AG action relies on the fact that its administration enhances butyric acid production in the bowel and hence rising its serum concentration [13].

Butyric acid is a short chain fatty acid (SCFA) that synthesized via the fermentation of otherwise non-digestible fiber by bacteria in the colon [48]. It has four actions; first, its raises IL 10 serum level [14, 49], second, it increases serum levels of IL-1 receptor antagonist (IL-1RA), third, it suppresses synthesis of transforming growth factor (TGF-*β*1) [13] and fourth, it fosters the expression of fetal hemoglobin in erythrocytes [26]. Each of the aforementioned actions has a direct effect on schistosomiasis infection outcome; IL10 regulates not just the intensity of egg-induced inflammatory responses, but also the coherence of granuloma structure, particularly deposition of collagen by fibroblasts around the periphery [50]. It also downregulates B7 MHC II, costimulatory molecules on APC [51], leading to hyporesponsive state through induction of T cells energy [52]. IL-1RA was reported before to cause *in vivo* depletion of exacerbated granuloma size and augmented regional cytokine production [53]. The effect of both IL 10 & IL-1RA might be manifested in this experiment in decreased granuloma diameters, fibrosis, increased cellularity and deteriorated *Schistosoma* eggs status inside the lesions.

Transforming growth factor (TGF-*β*1) is one of the strongest factors that lead to liver fibrosis. TGF-*β*1 promotes hepatic stellate cell (HSC) proliferation and collagen synthesis in the activated HSC [54] or modulates deposition of extracellular matrix (ECM) components and immune functions [55]. Furthermore, a number of researchers have recognized TGF-*β*1 inhibition as one of the factors that can be used to evaluate the antifibrotic effects of drugs on hosts infected with *Schistosoma japonicum* [56]. Consequently, possible suppression of TGF-*β*1 by AG could reverse the immunopathologic effect induced by *Schistosoma* eggs in the affected tissues as seen in in the current study.

Blood-feeding parasites, including schistosomes, hookworms, and malaria parasites, make use of aspartic proteases to produce initial or early cleavages in ingested host hemoglobin. Although phylogenetically distinct, these parasites all have the same food source; they are obligate blood feeders, or hematophagous. Hb from ingested or parasitized erythrocytes is their major source of exogenous amino acids for growth, development, and reproduction; the Hb, a 64-kDa tetrameric polypeptide, is broadly catabolized by parasite enzymes to free amino acids or small peptides [57].

The fact that fetal hemoglobin has been shown to slowdown hemoglobin degradation depriving *Schistosoma* worms of its food source [58], has inspired many researchers to evaluate the effect of increasing its production on murine malaria parasitemia [26], they reported that the administration of Arabic gum significantly decreased the parasitaemia and extended the lifespan of infected mice.

The present study demonstrated that AG was highly effective against the immature form of *S. mansoni* which resists PZQ, and using both agents together yielded the best results owing to their synergetic effect.

To recapitulate, the study in hands shed the light on a novel and “green” management approach of *Schistosomiasis mansoni*, being one of the safest dietary fibers, and perceptibly effective in treating immature forms which entails the abortion of reinfection in endemic areas. Further studies are on a larger scale are required to evaluate the feasibility of using AG as an effective treatment of immature schistosomiasis *mansoni* and for prophylaxis against reinfection, particularly in endemic areas where the control programs are continually hampered by many socioeconomic, topographic and cultural obstacles that are not currently anticipated to be defeated in the near future.

## References

1. Steinmann P, Keiser J, Bos R, Tanner M, Utzinger J. Schistosomiasis and water resources development: systematic review, meta-analysis, and estimates of people at risk. The Lancet infectious diseases. 2006;6(7):411-25.

2. King CH, Dangerfield-Cha M. The unacknowledged impact of chronic schistosomiasis. Chronic illness. 2008;4(1):65-79.

3. Taylor M. Global trends in schistosomiasis control. Bulletin of the World Health Organization. 2008;86(10):738-.

4. Fallon PG, Doenhoff MJ. Drug-resistant schistosomiasis: resistance to praziquantel and oxamniquine induced in Schistosoma mansoni in mice is drug specific. The American journal of tropical medicine and hygiene. 1994;51(1):83-8.

5. Ismail M, Botros S, Metwally A, William S, Farghally A, Tao L-F, et al. Resistance to praziquantel: direct evidence from Schistosoma mansoni isolated from Egyptian villagers. The American journal of tropical medicine and hygiene. 1999;60(6):932-5.

6. Ribeiro-dos-Santos G, Verjovski-Almeida S, Leite LC. Schistosomiasis—a century searching for chemotherapeutic drugs. Parasitology research. 2006;99(5):505.

7. Wang W, Wang L, Liang Y-S. Susceptibility or resistance of praziquantel in human schistosomiasis: a review. Parasitology research. 2012;111(5):1871-7.

8. Doenhoff MJ, Cioli D, Utzinger J. Praziquantel: mechanisms of action, resistance and new derivatives for schistosomiasis. Current opinion in infectious diseases. 2008;21(6):659-67.

9. Botros S, William S, Hammam O, Holý A. Activity of 9-(S)-[3-hydroxy-2- (phosphonomethoxy) propyl] adenine against Schistosomiasis mansoni in mice. Antimicrobial agents and chemotherapy. 2003;47(12):3853-8.

10. Rehan A, Johnson KJ, Kunkel RG, Wiggins RC. Role of oxygen radicals in phorbol myristate acetate-induced glomerular injury. Kidney international. 1985;27(3):503-11.

11. Al-Majed AA, Mostafa AM, Al-Rikabi AC, Al-Shabanah OA. Protective effects of oral arabic gum administration on gentamicin-induced nephrotoxicity in rats. Pharmacological Research. 2002;46(5):445-51.

12. Anderson D. Evidence for the safety of gum arabic (Acacia senegal (L.) Willd.) as a food additive—a brief review. Food Additives & Contaminants. 1986;3(3):225-30.

13. Matsumoto N, Riley S, Fraser D, Al-Assaf S, Ishimura E, Wolever T, et al. Butyrate modulates TGF-β1 generation and function: Potential renal benefit for Acacia (sen) SUPERGUM^TM^(gum arabic)? Kidney international. 2006;69(2):257-65.

14. Xuan NT, Shumilina E, Nasir O, Bobbala D, Götz F, Lang F. Stimulation of mouse dendritic cells by Gum Arabic. Cellular Physiology and Biochemistry. 2010;25(6):641-8.

15. Pradeep A, Happy D, Garg G. Short-term clinical effects of commercially available gel containing Acacia arabica: a randomized controlled clinical trial. Australian dental journal. 2010;55(1):65-9.

16. Gado AM, Aldahmash BA. Antioxidant effect of Arabic gum against mercuric chloride-induced nephrotoxicity. Drug design, development and therapy. 2013;7:1245.

17. Nasir O, Umbach AT, Rexhepaj R, Ackermann TF, Bhandaru M, Ebrahim A, et al. Effects of gum arabic (Acacia senegal) on renal function in diabetic mice. Kidney and Blood Pressure Research. 2012;35(5):365-72.

18. Nasir O. Renal and Extrarenal Effects of Gum Arabic (Acacia Senegal)-What Can be Learned from Animal Experiments? Kidney and Blood Pressure Research. 2013;37(4- 5):269-79.

19. Kaddam L, FdleAlmula I, Eisawi OA, Abdelrazig HA, Elnimeiri M, Lang F, et al. Gum Arabic as fetal hemoglobin inducing agent in sickle cell anemia; in vivo study. BMC hematology. 2015;15(1):19.

20. Al-Yahya AA, Asad M. Antiulcer activity of gum arabic and its interaction with antiulcer effect of ranitidine in rats. Biomedical Research. 2016;27(4).

21. Ahmed AA, Fedail JS, Musa HH, Musa TH, Sifaldin AZ. Gum Arabic supplementation improved antioxidant status and alters expression of oxidative stress gene in ovary of mice fed high fat diet. Middle East Fertility Society Journal. 2016;21(2):101-8.

22. Fedail JS, Ahmed AA, Musa HH, Ismail E, Sifaldin AZ, Musa TH. Gum arabic improves semen quality and oxidative stress capacity in alloxan induced diabetes rats. Asian Pacific Journal of Reproduction. 2016;5(5):434-41.

23. Clark D, Gazi M, Cox S, Eley B, Tinsley G. The effects of Acacia arabica gum on the in vitro growth and protease activities of periodontopathic bacteria. Journal of clinical periodontology. 1993;20(4):238-43.

24. Saini ML, Saini R, Roy S, Kumar A. Comparative pharmacognostical and antimicrobial studies of Acacia species (Mimosaceae). Journal of Medicinal Plants Research. 2008;2(12):378-86.

25. Singh B, Dubey S, Siddiqui M. Antimicrobial activity of natural edible gums. Journal of Pharmaceutical Sciences. 2015;3(11):2217-21.

26. Ballal A, Bobbala D, Qadri SM, Föller M, Kempe D, Nasir O, et al. Anti-malarial effect of gum arabic. Malaria journal. 2011;10(1):139.

27. Holanda J, Pellegrino J, Gazzinelli G. Infection of mice with cercariae and schistosomula of Schistosoma mansoni by intravenous and subcutaneous routes. Revista do Instituto de Medicina Tropical de Sao Paulo. 1974;16(3):132-4.

28. Nessim NG, Demerdash Z. Correlation between infection intensity, serum immunoglobulin profile, cellular immunity and the efficacy of treatment with praziquantel in murine schistosomiasis mansoni. Arzneimittelforschung. 2000;50(02):173-7.

29. Duvall RH, DeWitt WB. An improved perfusion technique for recovering adult schistosomes from laboratory animals. The American journal of tropical medicine and hygiene. 1967;16(4):483-6.

30. Abdel-Salam A, Ammar N, Abdel-Hamid A. Effectiveness of probiotic Labneh supplemented with garlic or onion oil against Schistosoma mansoni in infected mice. Int J Dairy Sci. 2008;3(2):97-104.

31. Cheever AW. Quantitative comparison of the intensity of Schistosoma mansoni infections in man and experimental animals. Transactions of the Royal Society of Tropical Medicine and Hygiene. 1969;63(6):781-95.

32. Selem RF, Eraky MA. Assessment of mefloquine in-vivo efficacy on juvenile and adult stages of Schistosoma haematobium (Egyptian strain). Parasitologists United Journal. 2015;8(1):60.

33. Cheever AW. Conditions affecting the accuracy of potassium hydroxide digestion techniques for counting Schistosoma mansoni eggs in tissues. Bulletin of the World Health Organization. 1968;39(2):328.

34. Pellegrino J, Oliveira CA, Faria J, Cunha AS. New approach to the screening of drugs in experimental schistosomiasis mansoni in mice. The American journal of tropical medicine and hygiene. 1962;11(2):201-15.

35. Cançado JR, da Cunha AS, de Carvalho DG, Cambraia JS. Evaluation of the treatment of human Schistosoma mansoni infection by the quantitative oogram technique. Bulletin of the World Health Organization. 1965;33(4):557.

36. Harris H. On the rapid conversion of haematoxylin into haematein in staining reactions. Journal of Applied Microscopic Laboratory Methods. 1900;3(3):777.

37. Masson P. Some histological methods: trichrome stainings and their preliminary technique. J Tech Methods. 1929;12:75-90.

38. Botros S, El-Badrawy N, Metwally A, Khayyal M. Study of some immunopharmacological properties of praziquantel in experimental schistosomiasis mansoni. Annals of Tropical Medicine & Parasitology. 1986;80(2):189-96.

39. Lichtenberg Fv. Host response to eggs of S. mansoni: I. Granuloma formation in the unsensitized laboratory mouse. The American Journal of Pathology. 1962;41(6):711.

40. Savioli L, Daumerie D. Sustaining the drive to overcome the global impact of neglected tropical diseases: second WHO report on neglected tropical diseases: World Health Organization; 2013.

41. Botelho MC, Oliveira PA, Vieira P, Delgado MdL, Lourenço L, Lopes C, et al. Granulomatous-like immune reaction and hepatic fibrosis induced by Schistosoma haematobium immature worms. Virulence. 2010;1(3):123-9.

42. Colley DG. Morbidity control of schistosomiasis by mass drug administration: how can we do it best and what will it take to move on to elimination? Tropical medicine and health. 2014;42(2SUPPLEMENT):S25-S32.

43. Ali BH, Ziada A, Blunden G. Biological effects of gum arabic: a review of some recent research. Food and Chemical Toxicology. 2009;47(1):1-8.

44. van Duivenvoorde LM, Han WG, Bakker AM, Louis-Plence P, Charbonnier L-M, Apparailly F, et al. Immunomodulatory dendritic cells inhibit Th1 responses and arthritis via different mechanisms. The Journal of Immunology. 2007;179(3):1506-15.

45. Ahmed AA, Fedail JS, Musa HH, Kamboh AA, Sifaldin AZ, Musa TH. Gum Arabic extracts protect against hepatic oxidative stress in alloxan induced diabetes in rats. Pathophysiology. 2015;22(4):189-94.

46. Al-Olayan EM, El-Khadragy MF, Alajmi RA, Othman MS, Bauomy AA, Ibrahim SR, et al. Ceratonia siliqua pod extract ameliorates Schistosoma mansoni-induced liver fibrosis and oxidative stress. BMC complementary and alternative medicine. 2016;16(1):434.

47. Soliman R, Ismail O, Badr M, Nasr S. Resveratrol ameliorates oxidative stress and organ dysfunction in Schistosoma mansoni infected mice. Experimental parasitology. 2017;174:52-8.

48. Pryde SE, Duncan SH, Hold GL, Stewart CS, Flint HJ. The microbiology of butyrate formation in the human colon. FEMS microbiology letters. 2002;217(2):133-9.

49. West NP, Christophersen CT, Pyne DB, Cripps AW, Conlon MA, Topping DL, et al. Butyrylated starch increases colonic butyrate concentration but has limited effects on immunity in healthy physically active individuals. Exercise immunology review. 2013;19.

50. Sadler CH, Rutitzky LI, Stadecker MJ, Wilson RA. IL-10 is crucial for the transition from acute to chronic disease state during infection of mice with Schistosoma mansoni. European journal of immunology. 2003;33(4):880-8.

51. Ding L, Linsley P, Huang L, Germain R, Shevach E. IL-10 inhibits macrophage costimulatory activity by selectively inhibiting the up-regulation of B7 expression. The Journal of Immunology. 1993;151(3):1224-34.

52. King CL, Medhat A, Malhotra I, Nafeh M, Helmy A, Khaudary J, et al. Cytokine control of parasite-specific anergy in human urinary schistosomiasis. IL-10 modulates lymphocyte reactivity. The Journal of Immunology. 1996;156(12):4715-21.

53. Ruth JH, Bienkowski M, Warmington KS, Lincoln PM, Kunkel SL, Chensue SW. IL-1 receptor antagonist (IL-1ra) expression, function, and cytokine-mediated regulation during mycobacterial and schistosomal antigen-elicited granuloma formation. The Journal of Immunology. 1996;156(7):2503-9.

54. Bowen T, Jenkins RH, Fraser DJ. MicroRNAs, transforming growth factor beta-1, and tissue fibrosis. The Journal of pathology. 2013;229(2):274-85.

55. Verrecchia F, Mauviel A. Transforming growth factor-β signaling through the Smad pathway: role in extracellular matrix gene expression and regulation. Journal of Investigative Dermatology. 2002;118(2):211-5.

56. Chen B-L, Zhang G-Y, Wang S-P, Li Q, Xu M-H, Shen Y-M, et al. The combined treatment of praziquantel with osteopontin immunoneutralization reduces liver damage in Schistosoma japonicum-infected mice. Parasitology. 2012;139(4):522-9.

57. Brinkworth RI, Prociv P, Loukas A, Brindley PJ. Hemoglobin-degrading, Aspartic Proteases of Blood-feeding Parasites SUBSTRATE SPECIFICITY REVEALED BY HOMOLOGY MODELS. Journal of Biological Chemistry. 2001;276(42):38844-51.

58. Shear HL, Grinberg L, Gilman J, Fabry ME, Stamatoyannopoulos G, Goldberg DE, et al. Transgenic mice expressing human fetal globin are protected from malaria by a novel mechanism. Blood. 1998;92(7):2520-6.

